# Fibroblast contractility drives network reorganization and epithelial proliferation in intestinal polyposis

**DOI:** 10.1101/2025.01.08.632004

**Authors:** Mei-Lan Li, Yuying Wang, Maria Figetakis, David Gonzalez, Jason Jin, Elizabeth S. McDonald, Nadia A. Ameen, Kaelyn Sumigray

## Abstract

Fibroblasts are critical regulators of epithelial homeostasis through mechanics and signaling. However, the regulatory principles governing fibroblast behaviors are largely unknown. Fibroblast dysregulation has emerged as a pathological contributor in epithelial diseases such as intestinal polyposis. Here, using an inducible Bmp-loss-of-function polyposis model, we define stepwise mechanisms unraveling how dysregulated signaling perturbs fibroblast behaviors, which in turn, disrupt their function to regulate epithelial homeostasis. Intriguingly, the first initiating event leading to epithelial polyps was architectural and impacted the fibroblasts, not the epithelium. Bmp signaling inhibition caused fibroblasts to become hypercontractile, leading to their reorganization from an elaborate network into collapsed clusters beneath crypts. Disrupted fibroblast mechanics and compartmentalization preceded epithelial hyperproliferation in polyps. Using in vitro models, we show that fibroblast hypercontractility not only disrupted fibroblast organization, but also enhanced the ability of fibroblasts to support epithelial growth, leading to epithelial dysregulation. Overall, our studies reveal stepwise regulatory mechanisms underlying fibroblast signaling, mechanics and organization critical for their function to regulate epithelial homeostasis.

## Introduction

Epithelial-mesenchymal crosstalk is a crucial mechanism regulating tissue homeostasis and architecture. Extensive efforts have identified heterogeneous populations of fibroblasts as critical niche players and revealed their secreted niche signals essential for epithelial homeostasis across organs^1–3^. Understand the fundamental principles governing epithelial-fibroblast crosstalk is critical, as dysregulated fibroblasts have pathogenic roles in tumorigenesis, inflammation and fibrosis^4–6^. The adult mammalian small intestine is a powerful system for studying epithelial-fibroblast crosstalk due to its rapid cellular turnover, unique ribbon-shaped tissue architecture, and well-characterized stem cell niches. Within the intestine, fibroblast subtypes compartmentalize and contribute to the opposing Wnt and Bmp signaling gradients that maintain stemness in crypts and promote epithelial differentiation in villi^7–13^. While we have some understanding about the function of fibroblasts in terms of ligand secretion, how fibroblasts are organized and regulated to maintain epithelial homeostasis and proper tissue architecture remain unknown.

Intestinal fibroblasts secrete Bmp signals, among other factors, to regulate epithelial differentiation^8,14^. Global loss of Bmp signaling leads to uncontrolled epithelial proliferation and massive, distorted tissue outgrowths characterized by polyps within the gastrointestinal tract. Two human polyposis syndromes, such as juvenile polyposis and hereditary mixed polyposis syndromes are caused by loss of function (LOF) in Bmp signaling, including germline mutations in the Bmp receptor *BMPR1A* or the downstream regulator *SMAD4*, and gene duplication in the Bmp antagonist *GREM1*^15–19^. As Bmp signals to epithelial stem cells for differentiation, it is largely thought that loss of epithelial Bmp activity permits epithelial overgrowth and polyp formation. However, genetic mouse models have demonstrated that specific loss of Bmp receptor from the intestinal epithelium is not sufficient to produce polyps^20^, whereas loss of Bmp receptor from fibroblasts resulted in gastrointestinal polyps^21–25^. This highlights the critical roles of fibroblasts in both regulating epithelial cell behaviors and maintaining proper tissue architecture. The mechanisms through which fibroblasts disrupt tissue homeostasis and drive polyp formation remain largely unknown.

Previous reports utilized constitutive overexpression of secreted Bmp antagonists Noggin or Grem1 to induce full-grown epithelial polyps that recapitulated human intestinal polyposis syndromes^26–28^. To define the cellular mechanisms in polyp initiation, we employed a mouse model of inducible Noggin expression in the intestine. Surprisingly, we find that the initial changes to cell behaviors and tissue architecture during polyp formation are within the niche, not the epithelium. This initial change in tissue architecture folds the intact epithelium away from its underlying submucosa in an ectopic internal tissue fold. Distinct fibroblast subpopulations normally segregated along the crypt-villus axis, but upon Bmp inhibition, become hypercontractile and cluster in ectopic tissue fold, prior to epithelial dysregulation. We then leverage primary intestinal fibroblast culture to demonstrate that fibroblast hypercontractility is the driving force behind the abnormal cell clustering from Bmp inhibition. Finally, we find that Noggin-induced fibroblasts enhance epithelial growth in intestinal organoids and that this ability is dependent on fibroblast hypercontractility. Together, our work reveals stepwise regulations of fibroblast homeostasis governing their signaling, mechanics and organization essential for their function to regulate epithelial homeostasis.

## Results

### Intestinal polyps initiate through architectural changes rather than epithelial hyperproliferation

Previous models of constitutive Noggin overexpression described de novo crypt formation and epithelial hyperproliferation with disrupted crypt-villus axes in polyps^26,27^. To define the initial cellular events during polyp formation, we utilized a doxycycline-inducible mouse model of Noggin overexpression (Fig S1A). Surprisingly, epithelial proliferation or deformation was not the first observable cellular change. Instead, within three days of Noggin overexpression, the first noticeable cellular change was within the stroma, where the tissue formed an ectopic fold (Fig 1A-A’). Notably, the epithelium along the ectopic fold contained intact crypt-villus axes indistinguishable from control and normal adjacent epithelium. The ectopic tissue fold separated the epithelium from its underlying stroma away from the smooth muscle wall into the intestinal lumen. This separation generated a substantial 2.3-fold increase in area of the submucosal stromal region below crypts (Fig 1B). Only after 1-2 weeks did ectopic tissue folds become replaced by polyps, hyperproliferative overgrowths of the epithelium with disrupted architecture and loss of crypt-villus axes (Fig 1A’’). Polyps, but not ectopic folds, often contained misaligned crypts orthogonal to the villus axis away from the underlying stroma (Fig S1B-B’’).

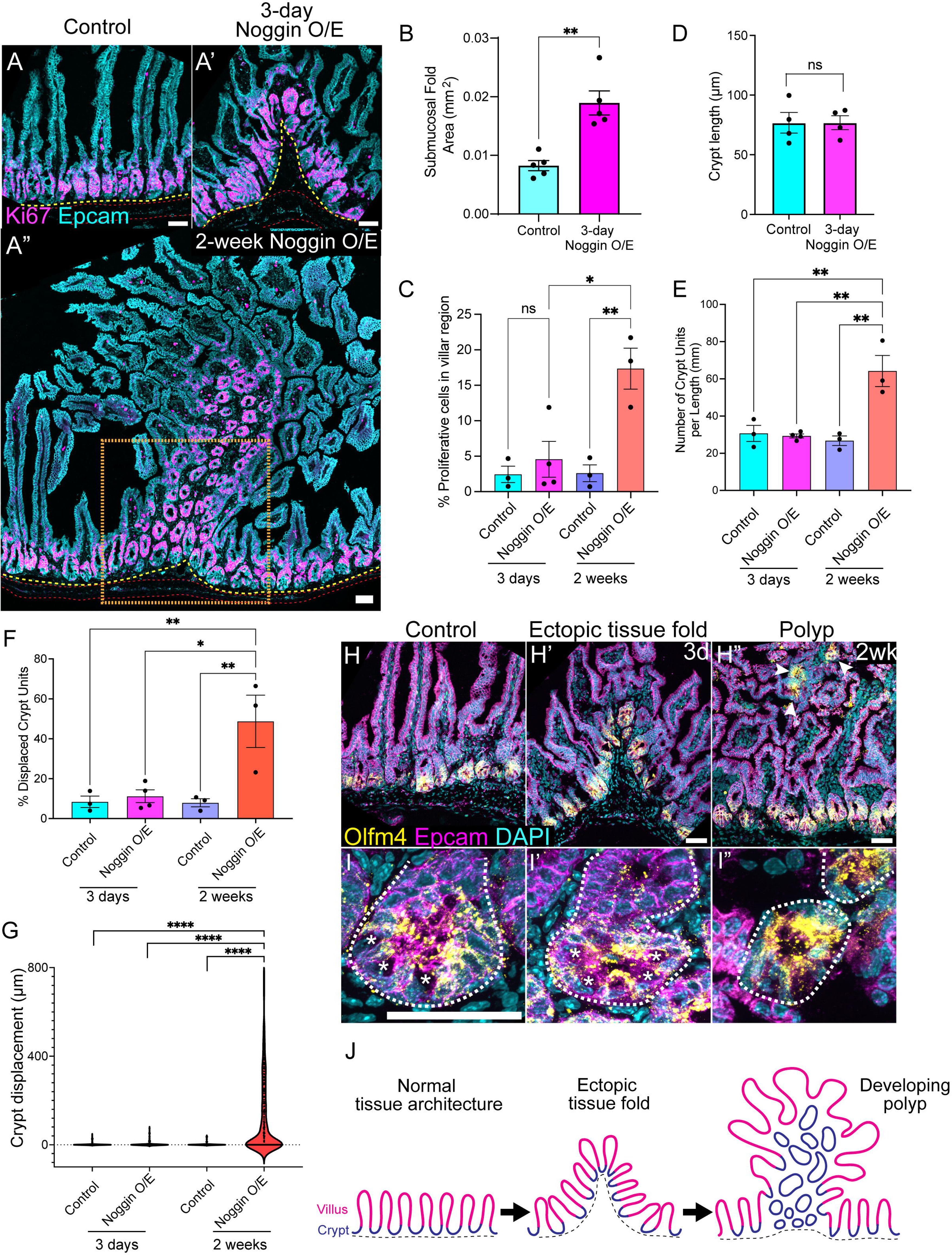

To define cellular and morphological changes to the epithelium, we first determined whether the epithelium within ectopic tissue folds still maintained proper compartmentalization, with proliferation restricted to the crypt domains. In ectopic tissue folds, Ki67+ epithelial proliferation was restricted to the crypt regions with proliferation rates and crypt lengths comparable to control tissue (Fig 1A-A’, C, D). In contrast, increased epithelial proliferation with disrupted crypt-villus axes was evident in long-term (2-week) Noggin overexpression, as demonstrated by a 6.7-fold increase in the relative abundance of Ki67+ epithelial cells within the villar region (Fig 1A’’, C), a 2.4-fold increase in number of crypts per gut length (Fig 1E), and crypt displacement into the villar region (Fig 1F). While all crypts within control and Noggin-overexpressing ectopic folds remained in their proper domain along the muscle wall in their stereotypic cup-like shape, long-term Noggin overexpression resulted in ectopic crypts, localized up to hundreds of microns away from their normal position during homeostasis (Fig 1G). Strikingly, displaced crypts within polyps contained Olfm4+ stem cells (Fig 1H-I), suggesting their ability to sustain long-term maintenance and support growth. Crypts in ectopic tissue folds, similar to controls, had the stereotypical pattern of Olfm4+ stem cells sandwiched between Paneth cells, which are distinguished by their large morphology and basally localized nuclei (Fig 1I-I’, asterisks). In contrast, displaced crypts within villar regions of polyps highly expressed Olfm4 without a clear pattern of cellular organization of Paneth cells, likely due to the misalignment and displacement of crypts within polyps (Fig 1I’’). Whether displaced crypts are truly formed de novo, as suggested by previous literature^26^, or whether they originate from the crypt region detached from the surrounding stroma as a long, convoluted and invaginated epithelium is still unclear.

Taken together, our data demonstrate that decreased Bmp signaling results in the formation of intestinal polyps: massive tissue outgrowths of hyperproliferative epithelium with disrupted crypt-villus axes. Surprisingly however, the first identifiable cellular changes during polyp formation are not to the epithelium. Instead, polyps are formed through an intermediate step in which the intact epithelium and its associate stroma buckle away from the muscle wall to generate an ectopic tissue fold (Fig 1J). How the fold transforms into a hyperproliferative polyp is unclear.

### Fibroblast architectural deformation precedes epithelial dysregulation in polyp formation

While epithelial proliferation and architecture remain unchanged during short-term Noggin overexpression, the ectopic tissue folds contained expanded submucosal stromal region below crypts (Fig 1B), prompting us to define specific alterations within the stroma. Fibroblasts, which are critical stem cell niche players, reside underneath the epithelium along the crypt-villus axis to regulate epithelial homeostasis^1,7,8,10,11,13^. Loss of Bmp signaling specifically within fibroblasts led to rapid polyp formation^21–23,25,29^ (Fig S2A-B’). The stem cell-supporting roles of fibroblasts and their polyp-initiating contribution from disrupted signaling thus led us to define specific changes within fibroblasts in ectopic tissue folds. As epithelial hyperproliferation is a characteristic feature of polyps, we first examined whether fibroblast proliferation was affected by decreased Bmp signaling. In Noggin-overexpressing intestines, fibroblasts did not become hyperproliferative, as assessed through quantification of Ki67+ PDGFRl1J-positive cells (Fig S2C). Therefore, epithelial hyperproliferation in polyps was likely not due to an increased number of fibroblasts secreting trophic factors.

As polyps are distinguished in part by disrupted epithelial compartmentalization, we investigated whether fibroblast compartmentalization and/or organization was disrupted in ectopic tissue folds. Intestinal fibroblasts can be broadly categorized into two main classes based on expression levels of the fibroblast marker PDGFRα, PDGFRα-low and PDGFRα-high, with distinct localization patterns and functions^8^. PDGFRα-low cells are primarily distributed in the submucosal region under crypts, while PDGFRα-high cells are enriched at the crypt-villus boundary and within villi (Fig 2A). Strikingly, fibroblasts form extensive protrusions to establish an elaborate interconnected network^30–32^ (Fig 2B). For ease in visualizing fibroblast distribution, we utilized the PDGFRα-H2B-GFP mouse line to track fibroblast nuclei. While overall fibroblast networks did not undergo gross organizational change in Noggin-overexpressing intestines (Fig 2C-C’), we surprisingly identified drastic local changes to fibroblast organization, specifically within ectopic tissue folds.

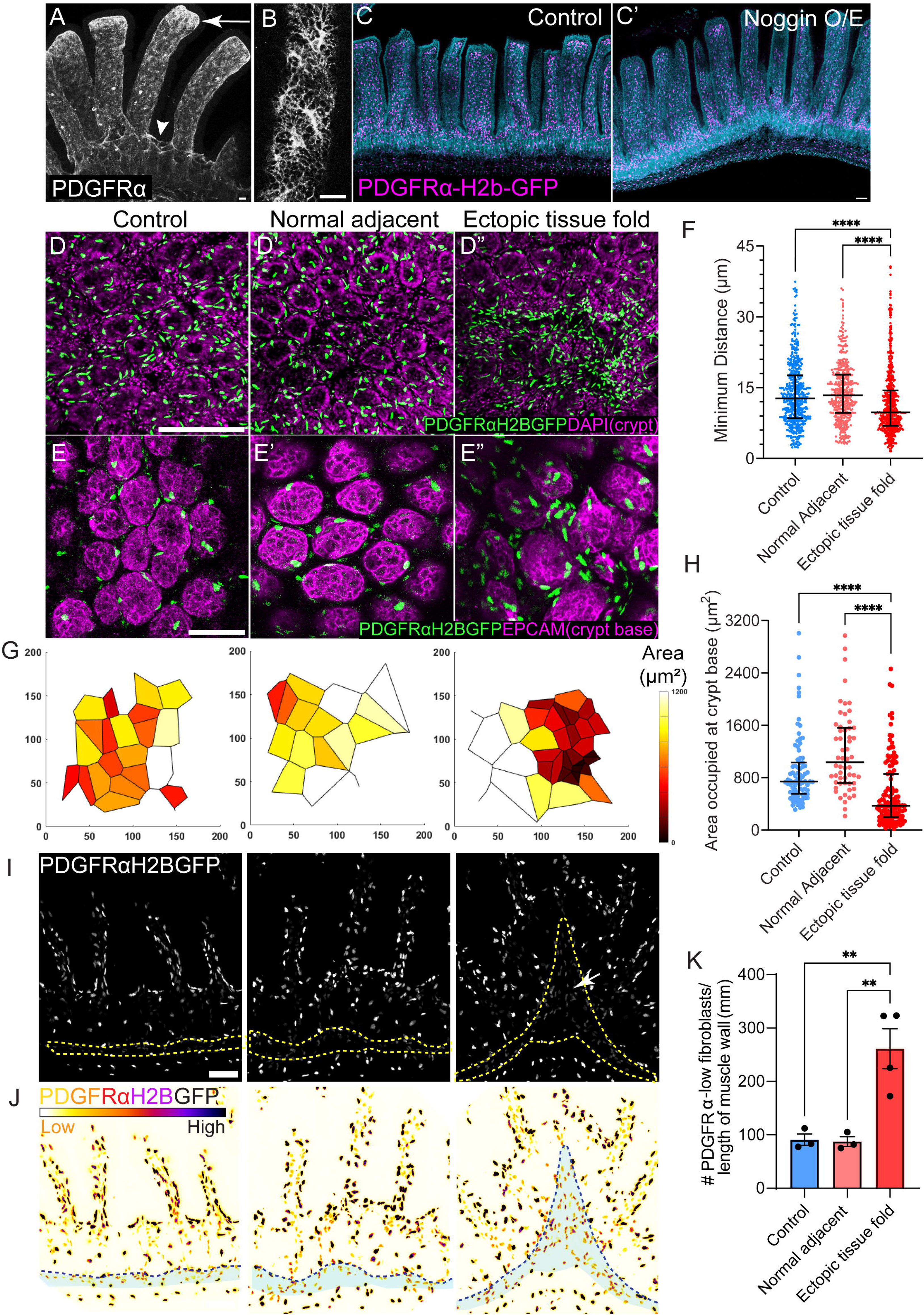

To examine fibroblast distribution in ectopic tissue folds, we imaged flat whole mount tissues through the muscle wall. This orientation allowed for visualization of crypts in cross-section and associated fibroblasts from the top-down view. Notably, PDGFRα-high fibroblasts rapidly reorganized and accumulated in ectopic tissue folds upon short-term Bmp inhibition (Fig 2D-D’’). In control and normal adjacent regions of Noggin-overexpressing intestines, PDGFRα-high fibroblasts were tightly associated with crypts, with their nuclei aligned along the circumference of each crypt (Fig 2E-E’). In contrast, PDGFRα-high fibroblasts in ectopic tissue folds were clustered beneath and around crypts and no longer aligned with crypts. Analysis of the 3D distance of each PDGFRα-high fibroblast with its nearest neighbor demonstrated decreased distances between cells, indicating increased local density and clustering of PDGFRα-high fibroblasts in ectopic tissue folds, whereas the distance between PDGFRα-high cells was unchanged in normal adjacent regions of Noggin-overexpressing intestines (Fig 2F). To analyze cell distribution specifically at crypt base, we utilized the PDGFRα-H2b-GFP nuclear signals to generate 2D Voronoi diagrams in the region of crypt bases to map PDGFRα-high cell distributions (Fig 2G). Voronoi diagrams created tessellation patterns based on each fibroblast and its closest neighbors to any other fibroblasts. A Voronoi cell polygon represented the area occupied by each PDGFRα-high fibroblast in the tessellation pattern based on distance calculation. Strikingly, fibroblasts in ectopic folds occupied less Voronoi areas than either control or normal adjacent regions, confirming the accumulation and clustering of PDGFRα-high fibroblasts in ectopic tissue folds, particularly at the crypt base region (Fig 2G-H). Together, these data demonstrate that PDGFRα-high fibroblasts mislocalize and cluster below crypts in ectopic tissue folds, distinct from their primary location in villi during homeostasis.

While PDGFRα-high cells comprise a distinct fibroblast subtype within the intestine, most intestinal fibroblasts express low levels of PDGFRα and are heterogeneous^8^. At least one subtype of PDGFRα-low fibroblasts is known to secrete trophic factors for epithelial growth, making the PDGFRα-low fibroblast population of particular interest in its potential to regulate epithelial homeostasis. To assess changes in the organization of PDGFRα-low fibroblasts within ectopic tissue folds, we focused on their distribution within the submucosal region below crypts (Fig 2I). Interestingly, PDGFRα-low fibroblasts became highly enriched within the expanded submucosal region of the ectopic tissue fold, while they maintained a distribution pattern similar to control animals and in normal adjacent regions of Noggin-overexpressing intestine (Fig 2J). Quantification of PDGFRα-low fibroblasts demonstrated a 2.9-fold increase in abundance below crypts within ectopic tissue folds (Fig 2K).

Taken together, our data demonstrate that distinct classes of fibroblast populations that are usually spatially segregated along the crypt-villus axis rapidly reorganize and cluster below crypts in ectopic tissue folds upon Bmp signaling inhibition. This disruption in fibroblast compartmentalization and architecture in ectopic tissue folds occurs prior to epithelial hyperproliferation in polyp initiation, demonstrating the relevance of fibroblast signaling and organization to maintain proper tissue homeostasis.

### Fibroblasts form a hypercontractile network in ectopic tissue folds

As intestinal fibroblasts are interconnected in networks, our surprising finding of fibroblast clustering in ectopic tissue folds prompted us to define the molecular and cellular behaviors behind this rapid reorganizational event. PDGFRα-high fibroblasts express low levels of myofibroblast markers^8,33–35^, and fibroblasts often transform into myofibroblasts in pathological conditions^36,37^, leading us to investigate the mechanical status of fibroblasts in ectopic tissue folds. While the expression level or pattern of myosin IIA and IIB expression level did not change in fibroblasts upon Noggin overexpression (data not shown), myosin activation was noticeably increased in fibroblasts within ectopic tissue folds, as shown by increased phosphorylated myosin light chain (pMLC19) staining (Fig 3A-B). Quantification of pMLC19 intensity in fibroblasts demonstrated a 2.5-fold increase in intensity (Fig 3C), suggesting that the fibroblast network around crypts becomes hypercontractile upon Bmp signaling inhibition. To further examine fibroblast contractility status, we stained intestines for Yap, a mechanosensitive transcriptional coactivator downstream of actomyosin contractility. As increased actomyosin contractility promotes nuclear translocation of Yap and subsequent activation^38^, we compared nuclear Yap intensity within PDGFRα-high and PDGFRα-low fibroblasts around and beneath crypts in control and ectopic tissue folds. Both classes of fibroblasts showed increased nuclear Yap signal (Fig 3D-F), further supporting the change in mechanical status of fibroblasts in Noggin-overexpressing intestines.

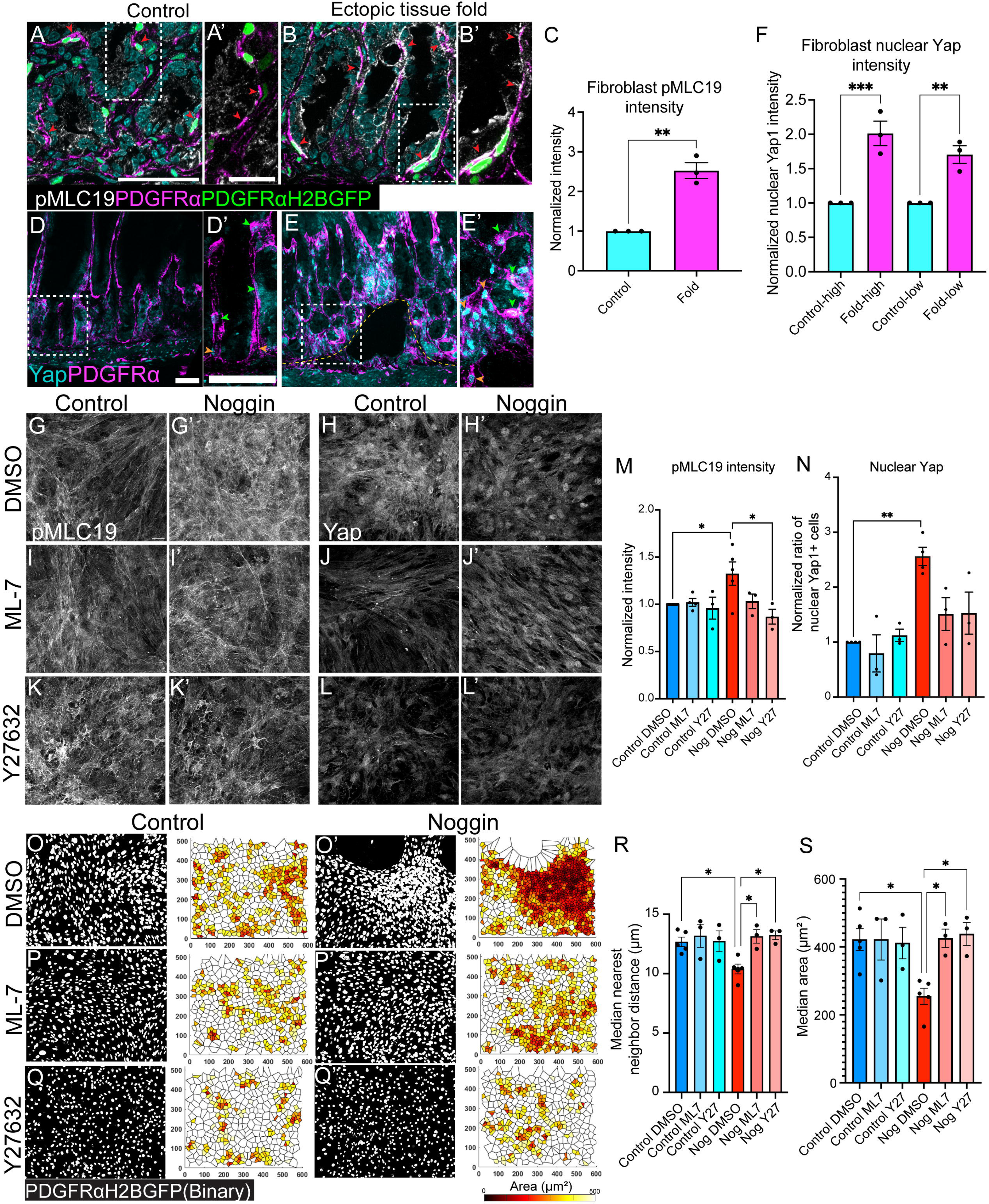

### Fibroblast contractility turns organized network into collapsed clusters

Taken together, our data support a model in which decreased Bmp signaling induces dramatic changes to fibroblast organization and mechanics prior to epithelial hyperproliferation during polyp initiation. The unique organization of fibroblasts as a continuous sheath underlying the epithelium and the increased fibroblast actomyosin contractility suggested possible coordinated movements within the fibroblast network, resulting in changes in cell organization from Bmp signaling inhibition. Therefore, we hypothesized that fibroblast hypercontractility induced fibroblast cell clustering. To test whether actomyosin contractility drove fibroblast clustering, we leveraged a primary adult intestinal fibroblast culture system with pharmacological contractility inhibitors to determine the direct effects of actomyosin contractility on cell organization. We first determined whether Noggin was sufficient to increase fibroblast contractility in vitro. Treatment of primary fibroblasts with 30% Noggin conditioned media did not alter myosin IIA or IIB levels (Fig S3A-B, G-H), but resulted in increased pMLC19 intensity (Fig 3G-G’, M) and nuclear Yap localization (Fig 3H-H’, N), consistent with our in vivo findings. Importantly, Noggin treatment also induced fibroblast clustering (Fig 3O-O’, R-S), reminiscent of the changes observed in vivo. Quantification of the distance between each cell and its nearest neighbor demonstrated that Noggin-treated fibroblasts were indeed closer within clusters (Fig 3R). Voronoi diagrams also showed that Noggin-induced cells occupied less Voronoi areas than control fibroblasts, confirming their clustering behavior (Fig 3O-O’, S).

In extreme cases, Noggin treatment turned uniformly distributed, confluent layers of fibroblasts into large aggregates connected to each other by bundles of elongated fibroblasts aligned in long cables (Fig S3I-J). Interestingly, these large fibroblast cables between cell clusters were sometimes “torn” or “recoiled”, suggesting possible pulling and coordinated cell movements driven by fibroblast hypercontractility. To test whether increased fibroblast contractility was responsible for cell clustering, we blocked actomyosin contractility using the myosin light chain kinase inhibitor ML-7 or ROCK inhibitor Y27632. Co-treatment of Noggin and either contractility inhibitor in primary fibroblasts returned contractility status back to control levels, as demonstrated by similar level of pMLC19 intensity and nuclear Yap localization (Fig 3G-N). Importantly, the co-treatment also rescued fibroblast cell clustering and cable formation, returning fibroblast organization from clustered to uniformly distributed cell network that resembled control fibroblasts (Fig 3O-Q). This rescue in cell organization was confirmed by distance measurements between nearest neighbors (Fig 3R) and Voronoi mapping between control and Noggin-treated fibroblasts in the presence of contractility inhibitors (Fig 3O-Q, S). Our in vitro study thus reveals that fibroblast hypercontractility drives cell clustering upon Bmp signaling inhibition, offering a potential mechanism underlying fibroblast reorganization observed in vivo.

### Fibroblast contractility is necessary for enhanced epithelial growth

Thus far, our data revealed critical regulatory roles of Bmp signaling in maintaining proper fibroblast mechanics, which in turn, preserves their unique organization. Next, we determined how dysregulated fibroblast Bmp signaling leads to epithelial overgrowth, the hallmark of intestinal polyps. To do this, we performed co-culture of primary intestinal fibroblasts and epithelial spheroids to model epithelial-fibroblast crosstalk in vitro. We first pre-treated primary fibroblasts with Noggin, then collected conditioned media to culture freshly isolated crypts in Matrigel for 4-6 days (Fig 4A). We assessed spheroid survival rate and luminal area as read-outs for growth. As fibroblasts are critical niche signal secretors that regulate epithelial behaviors, this experimental design allowed us to examine whether and how fibroblast Bmp inhibition affected their function in regulating epithelial growth, primarily through ligand secretion. Compared to media from control fibroblasts, conditioned media from Noggin-treated fibroblasts induced better and larger spheroid formation, as indicated by a 1.8-fold increase in spheroid luminal area (Fig 4B-C, F) and a 2.5-fold increase in survival (Fig 4G). These results reflected our in vivo observations of increased epithelial proliferation from inhibition of fibroblast Bmp signaling (Fig 1, S2A-B).

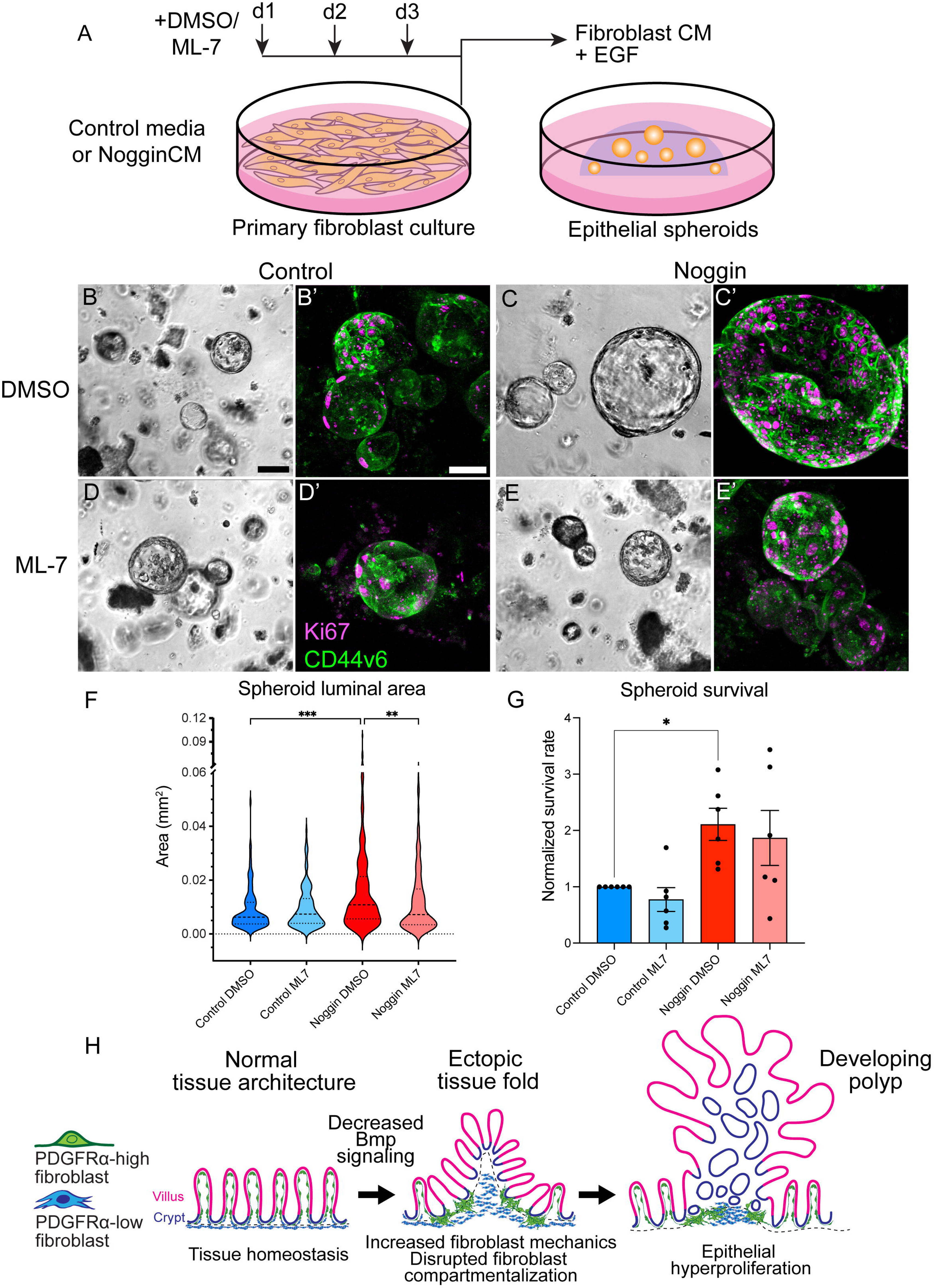

As increased fibroblast mechanics has pathological associations^36,37^, we hypothesized that fibroblast hypercontractility from Bmp inhibition was necessary to induce epithelial overgrowth. To test this, we pre-treated control or Noggin-induced fibroblasts with ML7 to rescue contractility from Noggin induction, then performed co-culture experiments with fibroblast conditioned media. Strikingly, fibroblast secretion from Noggin-ML7 co-treatment resulted in significantly decreased spheroid sizes relative to Noggin-treated fibroblasts, but comparable spheroid sizes to control fibroblasts (Fig 4B-F). Thus, contractility inhibition in Noggin-treated fibroblasts was sufficient to rescue organoid growth back to control levels. Interestingly, inhibition in fibroblast contractility did not alter spheroid survival count (Fig 4G). This suggests that fibroblast hypercontractility has a primary effect on promoting epithelial growth but not survival. The increased spheroid survival and growth observed in Noggin-treated fibroblasts could be mediated through separate mechanisms. Together, our data demonstrated that fibroblast hypercontractility from Bmp signaling inhibition enhances fibroblast function to support epithelial growth through differential ligand secretion.

## DISCUSSION

Overall, using intestinal polyp formation as a model, we defined regulatory roles of Bmp signaling in maintaining proper fibroblast mechanics and unique tissue architecture essential for their function in regulating epithelial proliferation (Fig 4H). During epithelial polyp initiation, the intestine first goes through an intermediate step involving changes in the niche architecture that results in ectopic tissue folding from decreased Bmp signaling. Fibroblasts, originally underlying the epithelium, become hypercontractile and cluster near and below crypts in ectopic tissue folds. These disruptions in fibroblast mechanics, organization and compartmentalization occur prior to epithelial hyperproliferation in polyp formation. Further mechanistic analyses using in vitro assays and pharmacological inhibitors suggest that fibroblast hypercontractility is a driving force of abnormal fibroblast clustering and epithelial hyperproliferation. Together, we propose that altered niche architecture synergized with impaired fibroblast mechanics and organization disrupts tissue homeostasis, resulting in epithelial dysregulation. As intestinal polyp is a consequence of dysregulated tissue homeostasis, the pathogenic niche-dependent mechanisms we identified contribute to the fundamental understanding of niche regulation critical for tissue homeostasis.

The continuous, ribbon-shaped intestinal epithelium compartmentalizes proliferation and differentiation into stem cell-containing crypts and absorptive villi. In intestinal polyps, this unique epithelial architecture is distorted into massive outgrowths with uncontrolled epithelial proliferation and disrupted crypt-villus axes. Interestingly, the first striking changes in tissue architecture and cell behaviors we observed in our Bmp-LOF polyposis mouse model are within the surrounding niche rather than the epithelium (Fig 1). This initial change in tissue architecture separates the normally shaped epithelium with intact crypt-villus axis from its surrounding niche, and expands the flat submucosal region below crypts into a prism-shaped space, resulting in an ectopic internal tissue fold. Distinct fibroblast subpopulations originally spatially segregated along crypt-villus axis are now mislocalized and accumulated below crypts in the ectopic tissue fold, indicating disrupted fibroblast compartmentalization and architecture (Fig 2). In tissue homeostasis, PDGFRα-low fibroblasts organized below crypts secrete R-spondin to promote stem cell proliferation^8^, whereas in ectopic tissue folds, they accumulate locally below crypts. It is possible that this fibroblast accumulation generates a concentrated niche signal gradient below crypts in ectopic tissue fold, thereby enabling subsequent epithelial hyperproliferation in a developing polyp. Similar ideas on how distinct tissue shape localizes niche signals to tightly control stem cell behaviors have been reported during villus and hair follicle morphogenesis^39^. In tissue homeostasis, PDGFRα-high fibroblasts localize along villi and accumulate at villus base, where they secrete Bmp to guide epithelial differentiation^8^. In ectopic tissue folds, these specialized fibroblasts are notably mislocalized from differentiated villi and clustered around proliferative crypts. This finding leads to questions on the role of fibroblast signaling and locational cue in specifying and maintaining fibroblast subtype identity and function during tissue homeostasis. Together our findings suggest that the initial change in tissue architecture creates a permissive environment that allows for subsequent epithelial deformation and hyperproliferation during polyp formation. These findings have broad implications for understanding how tissue architecture regulates tissue homeostasis.

Fibroblast hypercontractility and clustering are notable features of the ectopic tissue fold prior to epithelial hyperproliferation during polyp formation. Using in vitro primary fibroblast and organoid cultures, we show that fibroblast hypercontractility is necessary to drive fibroblast cell clustering and epithelial hyperproliferation upon Bmp inhibition (Fig 4). Recent studies indicated the roles of niche mechanics on tissue deformation and epithelial cell fate during tissue morphogenesis and homeostasis^40–42^. However, whether and how niche mechanics regulates epithelial proliferation in tissue homeostasis remained unknown. Our data here not only illustrate the role of fibroblast contractility in regulating epithelial proliferation, but also suggest differential ligand secretion as the mechanism underlying this mechanical regulation to support epithelial growth. This idea on altered fibroblast secretome aligns with a recent study that suggested changes in fibroblast secretome during polyp development^25^. It is exciting to reveal the connection between cell mechanics and ligand secretion in tissue homeostasis, as current efforts on cell mechanics focus on its roles in cell migration, fate specification and tissue deformation^42–45^, whereas studies on ligand secretion center on identification of niche players and their secreted signals in tissue homeostasis^1,7,8,10,11,13^. Our data here bridge together fibroblast mechanics and their functions in epithelial niche signaling, where hypercontractile fibroblasts drive epithelial hyperproliferation through differential ligand secretion.

In addition to epithelial dynamics, our data show that fibroblast mechanics also regulates their organization (Fig 2-3). In tissue homeostasis, fibroblasts are organized in networks underneath the epithelium to regulate epithelial cell behaviors^30–32^ (Fig 2B). Upon Bmp signaling inhibition, fibroblasts become hypercontractile and rapidly cluster beneath stem cells in vivo. Further analyses using small molecule inhibition in vitro show fibroblast hypercontractility as the driving force for cell clustering. Such mechanical-based regulatory control of cell organization was observed in tissue morphogenesis recently^40,46^, but not yet in tissue homeostasis or disease initiation. Together, our data here demonstrate the critical coordination of fibroblast signaling, mechanics and organization to perform proper function to regulate epithelial homeostasis, unraveling fundamental principles of fibroblast homeostasis and its contribution in disease initiation. These findings highlight how single cell behaviors sustain a global pattern of a multicellular tissue, opening foundational questions on the role of niche cell signaling, mechanics and organization in regulating tissue homeostasis.

The prevailing model of intestinal polyp formation attributes alterations in epithelial dynamics as the initiator. Studies have started to reveal the contribution of the stroma, in particular fibroblasts, in intestinal polyp formation using genetic knock-out of Bmp signaling receptor (Bmpr1a) to induce complete depletion of Bmp signaling in fibroblasts^21–23,25,29^ (Fig S2A-B’). Yet the rise of ectopic tissue folding as an initial local tissue architectural change prior to epithelial hyperproliferation has not been reported, nor the change in fibroblast mechanical status and disrupted fibroblast organization. It is possible that the lack of ectopic tissue folding in the published genetic Bmpr conditional knock-out models is due to difference in the dynamic range of Bmp signaling inhibition. Our inducible Bmp-LOF polyposis mouse model utilizes secreted protein Bmp antagonist Noggin, which sequesters Bmp outside of cell to prevent binding to its receptor to initiate Bmp signaling pathway intracellularly. This competitive signaling inhibition depends on the dynamic range of the secreted signal antagonist gradient, rather than a complete inhibition from the lack of signaling receptor. The rise of ectopic tissue folding in our system could be a result of a dampened Bmp signaling effect, rather than from a complete Bmp signaling loss. As an important lesson learned from developmental biology, the dynamic range of cell signaling is critical to orchestrate differential cell behaviors and organizations in tissue morphogenesis. Our findings have broad implications for understanding differential cellular response toward dynamic cell signaling in the context of disease initiation.

## Resource availability

### Lead contact

Requests for further information and resources should be directed to and will be fulfilled by the lead contact, Kaelyn Sumigray.

### Materials availability

This study did not generate new unique reagents.

### Data and code availability

All data reported in this paper will be shared by the lead contact upon request.

This paper does not report original code.

Any additional information required to reanalyze the data reported in this paper is available from the lead contact upon request.

## Acknowledgments

We thank all members of the Sumigray lab for thoughtful discussion and feedback and Dr. Valentina Greco for feedback and resource sharing. We also thank Dr. Mary Tomayko for reagents. This work was supported by the American Cancer Society awards IRG-17-172-57-IRG-03 and RSG-21-077-01-CCB (to K.S.) and NIH awards T32CA193200 and F31DK132866 (to M.L.L.).

## Author Contributions

Conceptualization, M.L and K.S.; Methodology: M.L, M.F., Y.W., D.G., and K.S.; Investigation, M.L., Y.W., M.F., J.J., E.S.M., and K.S.; Resources, N.A.A; Writing – Original Draft, M.L. and K.S.; Supervision, K.S.; Funding Acquisition, K.S.

## EXPERIMENTAL MODEL AND DETAILS

### Mice

All described animal work was approved by the Institutional Animal Care and Use Committee (IACUC) of Yale University. PDGFRα-H2BGFP (JAX strain #007669)^47^, Villin-rtTA*M2 (JAX strain #031283)^48^, Tre-Noggin-eGFP (JAX strain #023410)^49^ and PDGFRα-CreER (JAX strain #032770)^50^ were obtained from The Jackson Laboratory. Bmpr1a^fl/fl^ mice were kindly gifted by Dr. Yuji Mishina^51^. Adult mice older than 8 weeks, same sex-paired control and experimental littermates from both males and females were used for analysis. All mice were housed and maintained in a barrier facility with a 12h-12h light-dark cycle (07:00-19:00 light) and ad libitum access to food.

## METHOD DETAILS

### Tet-ON system and CreER induction

To induce Bmp signaling inhibition with the Tet-ON mouse system, Villin-rtTA; Tre-Noggin mice were on chow diet supplemented with 200 mg of doxycycline (Dox; VWR #89067-462) per kg of diet until harvest. Mice were harvested at various days after induction, 3 days to 2 weeks. Same-sex littermates of Villin-rtTA or Tre-Noggin control mice were on Dox chow together with mutant.To inhibit Bmp signaling with the conditional knock-out of Bmpr1a in fibroblasts, 5 mg tamoxifen (Sigma-Aldrich #T5648) in corn oil (Sigma #C8267) was administered via intraperitoneal injection into PDGFRα-CreER; Bmpr1a^fl/fl^ mice for five consecutive days. Same-sex littermates of Bmpr1a^fl/fl^ control mice were injected with the same tamoxifen scheme together with mutant.

### Tissue harvest

To prepare tissue cryosections, fresh mouse jejunum was embedded and frozen in optimal cutting temperature (OCT; Sakura Finetek USA #4583). Frozen OCT blocks were sectioned at 8 µm thickness using a cryostat.

To prepare whole mount tissues, mouse jejunum was first cut into 1 cm pieces, and then cut longitudinally to expose the epithelium. Flat intestinal pieces were pinned down on PDMS plates and fixed in 4% paraformaldehyde (PFA; Sigma-Aldrich #158127) at room temperature (RT) for two hours prior to overnight fixation at 4°C. After overnight fixation, whole mount tissues were washed with PBS three times before protection in sucrose gradient. Sucrose protection was done in the following order at RT: 5% sucrose in PBS for 5 min, 2:1 ratio of 5%:20% sucrose in PBS for 30 min, 1:1 ratio of 5%:20% sucrose in PBS for 30 min, 1:2 ratio of 5%:20% sucrose in PBS for 30 min, 20% sucrose in PBS overnight at 4°C. The next day, the sucrose protected whole mount tissues were stored in 20% sucrose in PBS containing 0.05% sodium azide. To prepare vibratome samples, mouse jejunum was cut into 2 cm pieces and fixed as tubes in 4% PFA at room temperature (RT) for two hours prior to overnight fixation at 4°C. Next day, fixed tissues were washed with PBS three times before storing in PBS containing 0.05% sodium azide. To create 300 µm vibratome sections, fixed tissues were embedded in 2.5% agarose (low-gelling temperature, Sigma-Aldrich #A9414), followed by cutting using Precisionary Compresstome (Model: VF-310-0Z) at speed 5.5 and oscillation 5.5 in PBS. Intestinal rings were stored in PBS at 4°C for up to two weeks prior to use.

### Primary fibroblast culture and treatment

Primary intestinal fibroblasts were isolated and cultured as described previously^52^. Briefly, longitudinally cut intestinal fragments in flat sheets were incubated in 20 mM EDTA in HBSS (Gibco #14170-112) for 20 min at 37°C. Fragments were then vortexed to deplete epithelial cells. After removal of muscle layers, tissues were finely minced and digested in enzyme mixtures of Collagenase II (1.5 mg/mL; Worthington Biochemical #LS004176) and Dispase II (1 mg/mL; Sigma-Aldrich #4942078001) in Complete DMEM media (Gibco #11995065) consisting of 10% FBS (Gibco #16-000-044) and 1x Pen-Strep (Gibco #15140-122) at 37°C on a nutating mixer for 10 min. Minced tissues were then vortexed to disrupt tissue clumps. After another 10 min of incubation, tissue suspension was passed through a 40 µm cell strainer to collect isolated mesenchymal cells on ice. Two more rounds of tissue digestion and cell collection were repeated in a total of 60 min incubation time. Cell suspension was treated with ACK buffer (Lonza #BP10-548E) for a minute on ice to remove red blood cells, then quenched with Complete DMEM. Isolated mesenchymal cells in the range of 0.47 million to 0.67 million cells/cm^2^ area were plated on glass coverslips for imaging and cultured in advanced DMEM/F12 media (Gibco #12634010) containing 10% FBS, 1X GlutaMAX (Gibco #35-050-061), 10 mM HEPES (Gibco #15630080) and 1X Pen-Strep with additional 1X Amphotericin B (Sigma-Aldrich #A2942) as anti-fungal. Amphotericin B was removed from culture after one day. Within 5-8 days in culture, fibroblasts attached and spread to reach confluence. To induce cell contractility, confluent primary fibroblasts were treated with Noggin conditioned media collected from 293FT-Noggin-Fc cells^53^ after dilution to 30% with base media Complete DMEM for three days. Control primary fibroblasts were treated with base media Complete DMEM simultaneously. To inhibit cell contractility, small molecule drug inhibitor ML-7 (10 µM; Cayman Chemical #11801) or Y-27632 (25 µM; Sigma-Aldrich #Y0503) was used in conjunction with Noggin for three days. Media were changed daily. Fibroblast conditioned media were collected at the end of day 3 of treatment for organoid culture.

### Organoid culture in fibroblast conditioned media

Fresh intestinal organoids were isolated as described previously^52^. Briefly, intestinal fragments were cut longitudinally to expose epithelium as flat sheets, then scraped with a glass coverslip to remove villi, followed by 30 min incubation in 3 mM EDTA in PBS on a rotator at 4°C. Remaining villi were released by shaking the epithelial sheets ice-cold PBS. When villi were mostly depleted, PBS was replaced and crypts were released by shaking the tissue in ice-cold PBS. Crypts were then collected, centrifuged (Thermo Scientific Megafuge 8) at 250 G for 5 min and washed once in ice-cold PBS before plating in 1X Matrigel (Corning #356231).

For organoid culture, fibroblast conditioned media were supplemented with recombinant mouse EGF protein (50 ng/mL; Thermo Fisher # PMG8041), 1X GlutaMAX, 10 mM HEPES, 1X B27 (Gibco #17504044), 1X N2 (Gibco #17502-048) and 1 mM N-acetyl-cysteine (Sigma-Aldrich #A9165). 10 µM Y-27632 and 1X Amphotericin B were used in fresh organoid culture and removed in subsequent media changes. Fibroblast conditioned media were replaced every other day in organoid culture.

### Immunofluorescence

For cryosection staining, sections were fixed for 8 min in 4% PFA, followed by a wash with PBS-T containing 0.2% Triton X-100 (AmericanBIO #AB02025). Sections were incubated in blocking buffer containing 3% BSA (Sigma-Aldrich #A9647) and 5% NDS (Jackson ImmunoResearch Labs #017-000-121) in PBS-T for at least 15 min. Then, primary antibodies diluted in blocking buffer were added to sections for 15 min at RT or overnight at 4°C. After PBS-T washing, secondary antibodies diluted in blocking buffer in 1:200 were applied onto sections for 10 min. After PBS-T washing, sections were mounted in antifade medium, consisting of 90% glycerol (JT Baker #2136-01) in PBS plus 2.5 mg/ml *p*-Phenylenediamine (Acros Organics #417481000).

For whole mount and vibratome section staining, tissues were placed in blocking buffer overnight at RT on a nutating mixer, followed by primary antibody incubation for 2-3 days at 37°C. Tissues were washed three times in PBS-T containing 1% Triton X-100 then incubated in secondary antibodies for 1-2 days at 37°C. Then, PBS-T washes were repeated prior to tissue clearing with Ce3D medium^54^ overnight at RT. Tissues were imaged in Ce3D clearing medium. For organoid staining, fixed organoids were first incubated in blocking buffer for 45 min at RT on a nutating mixer, then primary antibodies for 45 min at RT. After PBS-T washes, secondary antibodies were applied to organoids for 30 min at RT. Organoids were washed again with PBS-T before mounting and imaging in antifade solution on coverslips in a region contained by VALAP. For primary fibroblast monolayer staining, fixed cells on coverslips were incubated in PBS-T for 10 min, followed by the same blocking and staining steps as cryosection staining. Cells were mounted in ProLong glass antifade mountant (Invitrogen #P36980).

The following primary antibodies were used: Rabbit anti Ki67 (Abcam #ab15580), rat anti Epcam (Biolegend #118202), rabbit anti Olfm4 (Cell Signaling Technology #39141), rat anti CD44v6 (Thermo Fisher #BMS145), goat anti PDGFRα (R&D Systems #AF1062), rabbit anti MHCIIA (Biolegend #909801), rabbit anti MHCIIB (Biolegend #909902), rabbit anti phospho-MLC (S19) (Cell Signaling Technology #3671), rabbit anti phospho-MLC (T18/S19) (Cell Signaling Technology #3674), rabbit anti Yap (Cell Signaling #14074), and rabbit anti α-SMA (abcam #ab5694). Tissue sections and fibroblast monolayers were imaged on an upright Zeiss AxioImager with Apotome 2 attachment and Zeiss AxioCam 506 mono camera using Zen software (v3.0; Zeiss). Objectives used were Plan Apochromat 10x/0.45 air, 20x/0.8 air, 40x/1.3 oil, and 63x/1.4 oil. Whole mount tissues and vibratome sections were imaged on an inverted Leica Stellaris 5 confocal laser scanning microscope using Diode 405, a white light laser and LAS-X software (v4.6.1; Leica). Objectives used were HC PL APO CS2 10x/0.40 dry, 40x/1.10 water, 40x/1.30 oil, 63x/1.40 oil and HC FLUOTAR L VISIR 25x/0.95 water. Brightfield images of live organoid cultures were acquired on an upright Nikon Eclipse TS100 with camera QICAM Fast1394 using QCapture Pro software (v6.0; QImaging). Objectives used were Nikon Plan Fluor 4x/0.13 air and 10x/0.30 air.

## QUANTIFICATION AND STATISTICAL ANALYSIS

### Image analysis

The following quantifications were performed on Z-stacks of tissue cryosections or fibroblast monolayer cultures using Fiji/ImageJ (NIH): 1) Area of ectopic tissue fold, 2) epithelial proliferation, 3) number of crypts per length, 4) abundance of displaced crypts, 5) distance of crypt displacement, 6) number of accumulated PDGFRα-low fibroblasts, 7) fibroblast proliferation, 8) fibroblast contractility at ectopic tissue fold, 9) nuclear Yap intensity in fibroblasts at ectopic tissue fold, and 10) fibroblast contractility in cultured primary fibroblasts. Quantifications of 1-6 were done on images taken from the lateral view of tissues that captured crypt-villus axes. Spheroid survival rate was calculated as the number of live spheroids in treatment groups divided by the number of live spheroids in control. Luminal area of spheroids was measured by contouring the outer boundary of spheroids in focused brightfield images using Fiji/ImageJ.

Area of the ectopic tissue fold measured the expanded region between crypt base and muscle layers. 2) Epithelial proliferation was measured by first drawing selections to define crypt and villus region based on control, then quantifying the number of Ki67+ epithelial cells within each region relative to control. 3) Crypt number per length was measured by quantifying the number of crypts along the length of crypt base. 4) Abundance of displaced crypts was measured by first drawing a selection line to define muscularis mucosa, then counting Olfm4+ crypts that are not touching the line selection. 5) Distance of displaced crypts was quantified by first drawing a line selection to indicate crypt base, then drawing a perpendicular line from Olfm4+ crypts to the line selection to indicate distance away from crypt base. 6) Number of PDGFRα-low fibroblasts within submucosal area below crypts was quantified by first outlining the region between crypt base and muscle layers, then counting the number of PDGFRα-low fibroblasts within this region. 7) Fibroblast proliferation was measured based on the percentage of Ki67+ fibroblasts. 8) Fibroblast contractility at ectopic tissue fold was measured based on the intensity of phospho-MLC (S19) within PDGFRα+ fibroblast projections at 63x for five single z-slices per image. At least five images were quantified per animal. 9) Fibroblast Yap signaling activity was quantified based on intensity of Yap staining within fibroblast nuclei in region below crypt top/hinge and above muscle layers. 10) In vitro fibroblast contractility was measured based on intensity of MHCIIA, MHCIIB or phospho-MLC (S19) signal divided by intensity of control. Relative yap signaling activity in cultured fibroblasts was quantified based on intensity of Yap staining within fibroblast nuclei divided by nuclear intensity of control.

To measure fibroblast cell distribution, in vivo and in vitro quantifications of minimum distance and Voronoi area on fibroblasts were performed using image analysis software Aivia (v14.0.0; Leica) and Imaris (v10.1.1; Bitplane) along with a MATLAB coding script. 42 µm Z-stacks of whole mount images from crypt base to region below crypt top/hinge were first visualized and segmented in Aivia. Then, the spatial coordinates of PDGFRα-H2BGFP-high fibroblasts were extracted based on centroid positions and computed for minimum centroid distance. Minimum centroid distance was defined as the distance of a cell to its closest cell neighbor in 3D space, using the Euclidean distance formula. A fibroblast in clusters had shorter distance from its nearest neighbor than uniformly distributed fibroblasts. Voronoi diagrams were computed using the x, y coordinates of PDGFRα-H2BGFP-high fibroblasts within 12 µm z-stacks of whole mount images from crypt base, using a modified MATLAB coding script^55^. The presented Voronoi diagrams created tessellation polygon patterns based on each PDGFRα-high fibroblast and its neighboring cells closer to the cell than to any other fibroblasts. A Voronoi cell region represented the area occupied by each fibroblast in the tessellation pattern based on Euclidean distance. A fibroblast in clusters occupied less area in Voronoi diagrams than uniformly distributed fibroblasts. This analysis pipeline was also applied for 2D fibroblast distribution measurements in primary fibroblast monolayers. Median measurements of minimum distance and Voronoi cell area were plotted for the skewed data distributions in fibroblast monolayer culture.

### Statistics and reproducibility

Statistical calculations were performed with Prism 10 (GraphPad). Statistical parameters including the exact value of n, the definition of center, dispersion and precision measures (mean ± SEM or median with interquartile range) and statistical significance were reported in the Figure Legends. Data were considered to be statistically significant when *p* < 0.05. Asterisks in figures represented statistical significance calculated using the following statistical methods (NS = not significant; ^∗^, *p* < 0.05; ^∗∗^, *p* < 0.01; ^∗∗∗^, *p* < 0.001; ^∗∗∗∗^, *p* < 0.0001). Unpaired two-tailed t-test was performed for two comparisons. When multiple comparisons were made, one-way ANOVA with Tukey’s multiple comparisons test was used for normally distributed data. For skewed data distribution, the Kruskal-Wallis test with Dunn’s multiple comparisons test was performed instead. All experiments were repeated in at least three independent biological replicates.

---

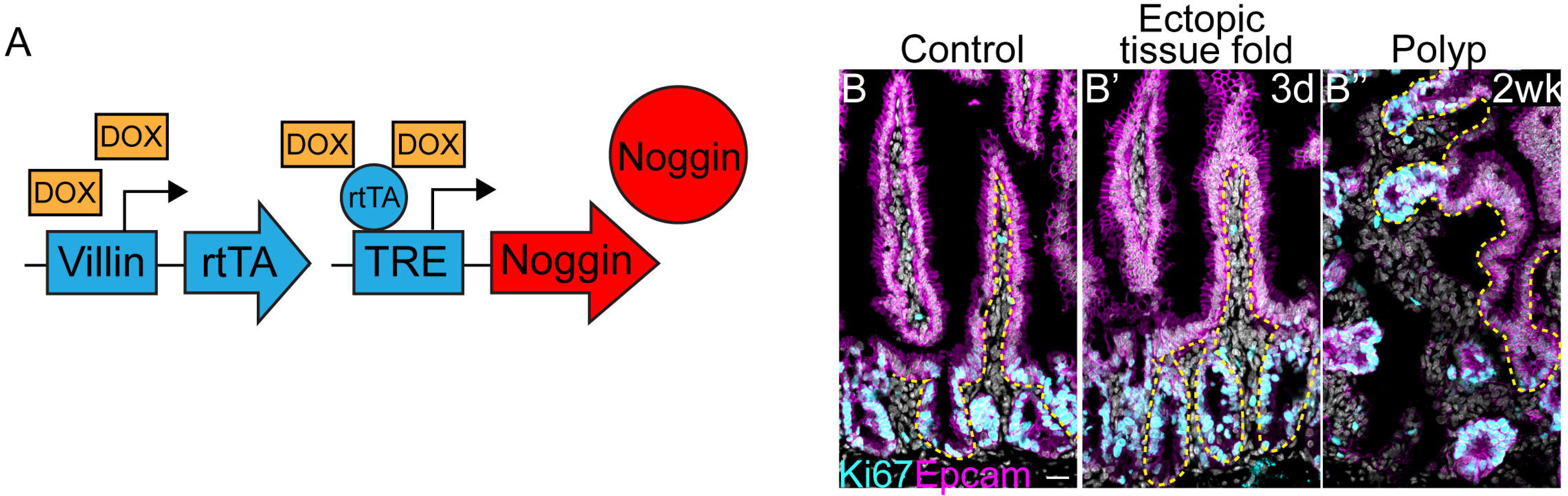

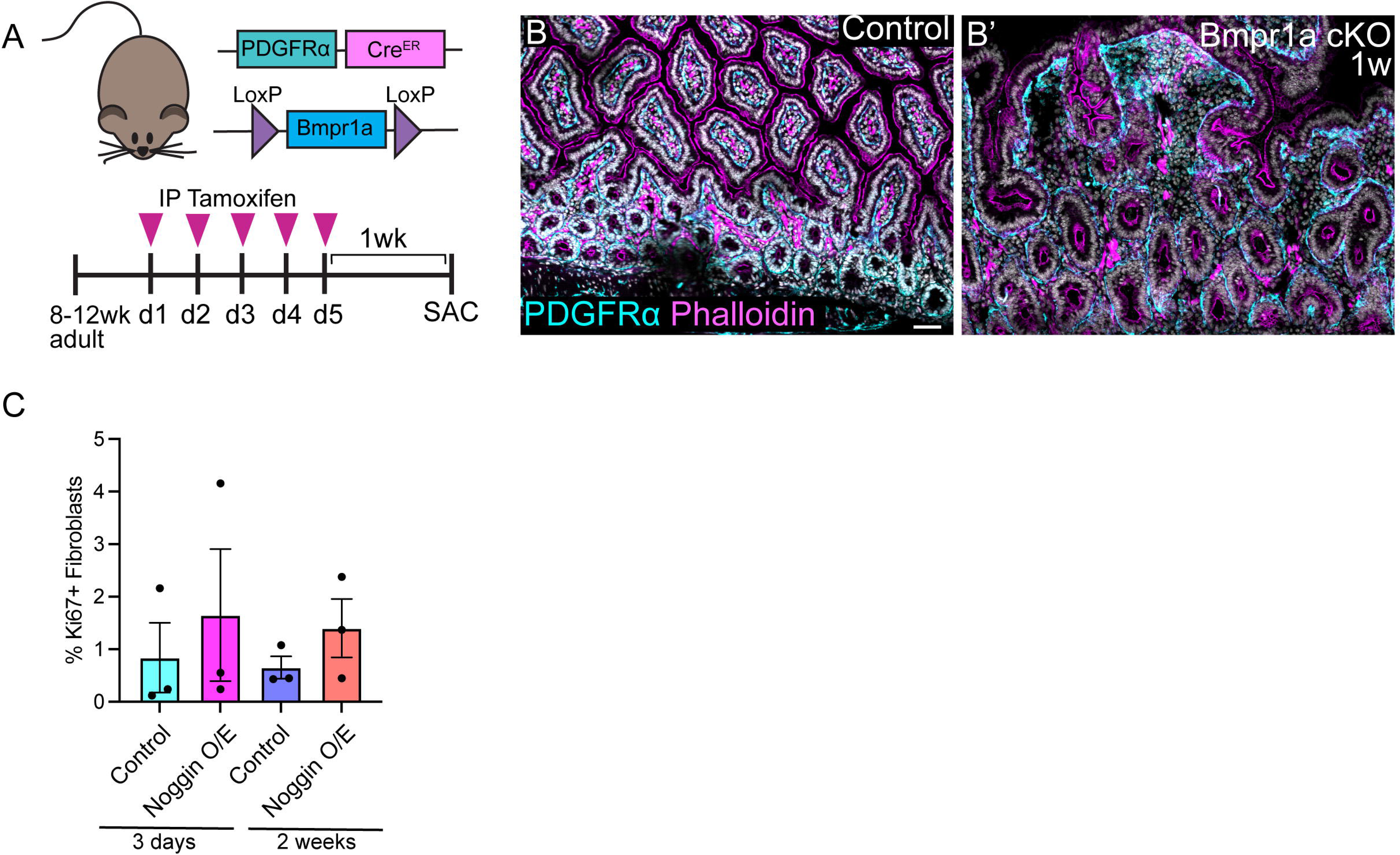

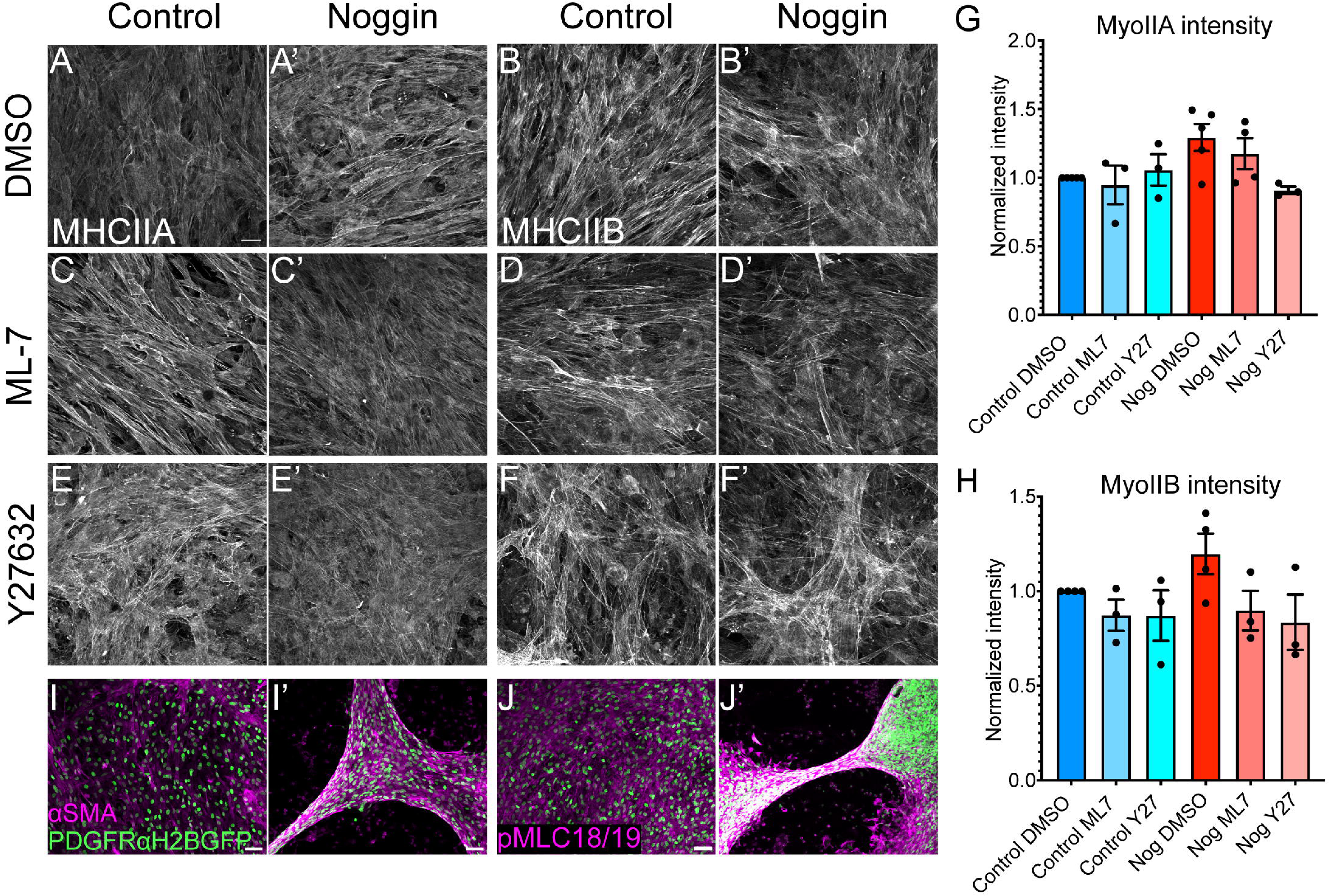

